# Dominant negative mutations in yeast Hsp90 reveal triage decision mechanism targeting client proteins for degradation

**DOI:** 10.1101/2024.01.02.573950

**Authors:** Julia M. Flynn, Margot E. Joyce, Daniel N. A. Bolon

**Affiliations:** Department of Biochemistry and Molecular Biotechnology University of Massachusetts Chan Medical School Worcester, MA 01605 USA

## Abstract

Most of the fundamental processes of cells are mediated by proteins. However, the biologically-relevant mechanism of most proteins are poorly understood. Dominant negative mutations have provided a valuable tool for investigating protein mechanisms but can be difficult to isolate because of their toxic effects. We used a mutational scanning approach to identify dominant negative mutations in yeast Hsp90. Hsp90 is a chaperone that forms dynamic complexes with many co-chaperones and client proteins. In vitro analyses have elucidated some key biochemical states and structures of Hsp90, co-chaperones, and clients; however, the biological mechanism of Hsp90 remains unclear. For example, high throughput studies have found that many E3 ubiquitin ligases bind to Hsp90, but it is unclear if these are primarily clients or acting to tag other clients for degradation. We introduced a library of all point mutations in the ATPase domain of Hsp90 into yeast and noticed that 176 were more than 10-fold depleted at the earliest point that we could analyze. There were two hot spot regions of the depleted mutations that were located at the hinges of a loop that closes over ATP. We quantified the dominant negative growth effects of mutations in the hinge regions using a library of mutations driven by an inducible promoter. We analyzed individual dominant negative mutations in detail and found that addition of the E33A mutation that prevents ATP hydrolysis by Hsp90 abrogated the dominant negative phenotype. Pull-down experiments did not reveal any stable binding partners, indicating that the dominant effects were mediated by dynamic complexes. DN Hsp90 decreased the expression level of two model Hsp90 clients, glucocorticoid receptor (GR) and v-src kinase. Using MG132, we found that GR was rapidly destabilized in a proteasome-dependent fashion. These findings provide evidence that the binding of E3 ligases to Hsp90 may serve a quality control function fundamental to eukaryotes.

## Introduction

Understanding protein mechanism is fundamental to understanding life because proteins carry out most of the biochemical reactions in living organisms. Where protein mechanisms have been successfully revealed, they have led to key biological insights^1,2^ and valuable biotechnology^3^. While science has made tremendous strides in illuminating ground state structures of many proteins, protein function often relies on transiently populated states (e.g. transition states in enzymes or dynamic binding interactions) that remain challenging to analyze structurally. Investigating the mechanism of proteins that function as interaction hubs is made more difficult because of the importance of studying function with all relevant binding partners present.

Dominant negative mutations that impede function in the presence of a wildtype (WT) copy provide a powerful approach to identify key mechanistic steps^4,5^. Dominant negative mutations have a rich history in the study of protein mechanism, though they have not been widely utilized in part because they are difficult to identify by traditional genetic screens. We recently developed mutational scanning approaches to identify dominant negative mutations in yeast^6,7^. In these studies, systematic libraries of point mutations in a gene were expressed and toxic variants that slowed growth in a bulk competition were identified based on decreased frequency using a deep sequencing readout. Here, we used mutational scanning to study dominant negative mutations in the Hsp90 molecular chaperone.

Hsp90 is an essential chaperone in eukaryotes^8^ where it is required for the maturation of client proteins to mature and active conformations^9^. While many chaperones including Hsp40 and Hsp70 bind to unfolded or misfolded conformations, Hsp90 binds to many clients in folded conformations that closely resemble their structure in the absence of chaperone^9^. High-throughput studies indicate that more than half of human kinases bind to Hsp90^10^. High-throughput analyses also show that about 30% of mammalian E3 ligases bind to Hsp90^10^, though it is not clear how many of these are clients of Hsp90 and/or act to stimulate degradation of Hsp90 clients. Hsp90 also assists in ligand binding to many nuclear steroid hormone receptors^11^. For example, studies of glucocorticoid receptor (GR) show that the steroid binding site is located in the interior of the protein structure and inaccessible to ligand binding in the absence of Hsp90^12,13^.

Small molecule inhibitors of Hsp90 have shown promise as anti-cancer agents^14^. Early studies demonstrated that geldanamycin, a natural product known to revert the transforming phenotype of v-src in mammalian cells in culture acted indirectly by inhibiting Hsp90 rather than v-src^15^. Potent synthetic inhibitors of Hsp90 have been developed and investigated as anti-cancer therapeutics^16^. While Hsp90 inhibitors have not yet been approved for therapeutic use, they have revealed important insights into Hsp90 mechanism. For example, Hsp90 inhibitors stimulate the ubiquitin-dependent degradation of many Hsp90 client proteins^17^. The association of many human E3 ubiquitin ligases with Hsp90^10^ suggests that some of these E3 enzymes function with Hsp90 in triage decisions where clients that cannot be matured to active conformations are instead targeted for degradation. Strong evidence shows that the human E3 CHIP serves this triage function with Hsp90. CHIP contains a TPR domain that binds in its native state to the C-terminus of Hsp90 leaving its ubiquitin ligase domain proximity to clients^18–20^. While the triage role of CHIP is known, the potential role of other E3’s in triage decisions is unclear. Of note, there is no clear homologue of the CHIP E3 ligase in yeast, and it is unknown if the triage function of Hsp90 is conserved across eukaryotes. It is also unclear how Hsp90 functions with CHIP or other E3’s to make triage decisions. For example, what are the molecular signatures that determine if a client will be chaperoned or targeted for destruction?

Hsp90 is a homodimeric ATPase and great progress has been made in understanding the ATPase driven conformational changes of this chaperone^21,22^. Hsp90 contains three domains referred to as N-terminal (N), Middle (M), and C-terminal (C). The C domain forms a stable homodimer ^23^, while the N domain is the site of nucleotide binding and can form a transient dimer that is stimulated by ATP, but not ADP binding^24,25^. In full length Hsp90, dimerization of the N domain leads to a closed state while N-domain dissociation leads to an open state. Large structural transition between the open and closed states are caused primarily by linker regions between the N, M, and C domains. In addition to the changes in the inter-domain linker regions, the other major ATP-driven conformation of Hsp90 is in the lid region of the N domain that closes over ATP, but not ADP or nucleotide-free structures^26^. How these ATPase driven conformational changes lead to client maturation remain unclear^27^ and are a major focus of current research efforts.

We investigated dominant negative mutations as a new approach to study mechanism in Hsp90. Based on current knowledge of Hsp90, we anticipated three main categories of dominant negative Hsp90 mutations (Figure 1). Cross dimerization of a mutant subunit with WT Hsp90 may lead to a defective heterodimer for many reasons (e.g. loss of binding to critical client or co-chaperone), that we reasoned would be difficult to elucidate in detail. To minimize this dominant mechanism in our analyses, we used an engineering strategy that prevents WT from cross dimerizing with our mutant variants (further described in results and methods). Based on previous consideration of dominant negative mutations^4^, we also anticipated that they may lead to tight binding to a critical partner protein making it inaccessible to WT Hsp90 (Figure 1B). Based on Hsp90’s role with CHIP in triage decisions, we also speculated a dominant mechanism where variants of Hsp90 stimulated the degradation of critical client proteins (Figure 1C).

**Figure 1.**
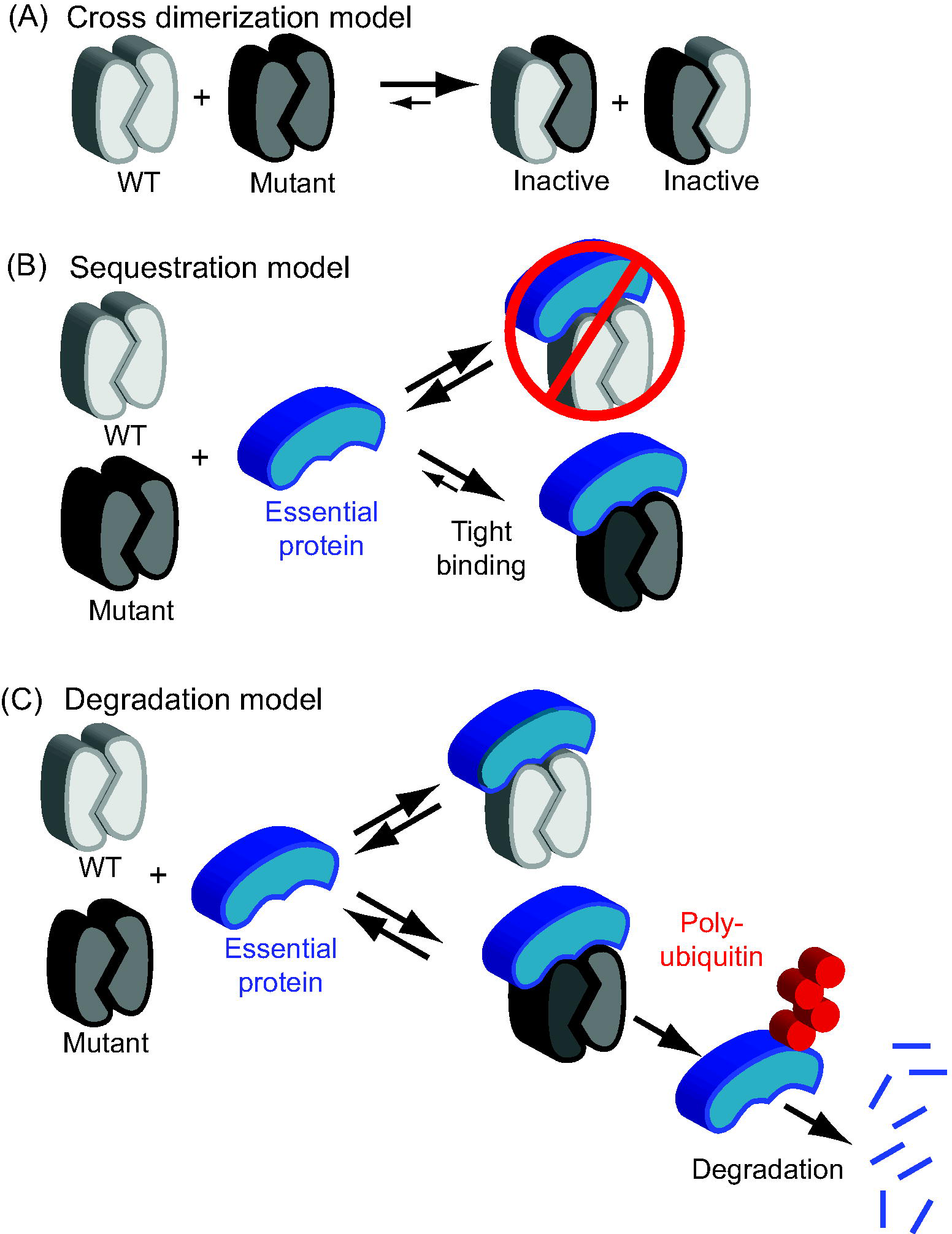
Potential dominant negative mechanisms for Hsp90. (A) Formation of inactive heterodimers with WT subunits. (B) Tight binding to critical binding partners that sequesters them from interacting with WT Hsp90. (C) Stimulation of the degradation of essential proteins.

## Results and Discussion

Our initial interest in dominant variants came from analyses of libraries of mutations in the ATPase domain of Hsp90 after they were transformed into yeast. Our initial goal was to assess the effects of these mutations as the sole source of Hsp90 in yeast. We utilized an engineered yeast strain where the wild-type copy of Hsp90 was driven by a regulatable promoter that could be turned on or off. After transforming libraries of Hsp90 variants into the engineered strain, cells that took up plasmid were selected for two days with the wild-type copy “on” at which point a sample of the culture was analyzed (Fig. 2A). When the expression of the mutant Hsp90 variants was driven by a promoter that mimics the endogenous expression of Hsp90^28^, we noticed that many mutations were depleted more than 10-fold during plasmid selection (Fig. 2B and Table S1). Depletion of these variants was impacted by promoter strength. In our previous work where libraries of Hsp90 mutants were driven by a promoter that results in roughly two orders of magnitude less Hsp90 protein^29^, almost all mutations were readily observed after plasmid selection. The same yeast strain and experimental approaches were used for analyzing mutants, indicating that the differences in depletion after plasmid selection were not due to detection or sampling limitations. Instead, these observations suggest that depletion was due to expression of toxic variants at high levels.

**Figure 2.**
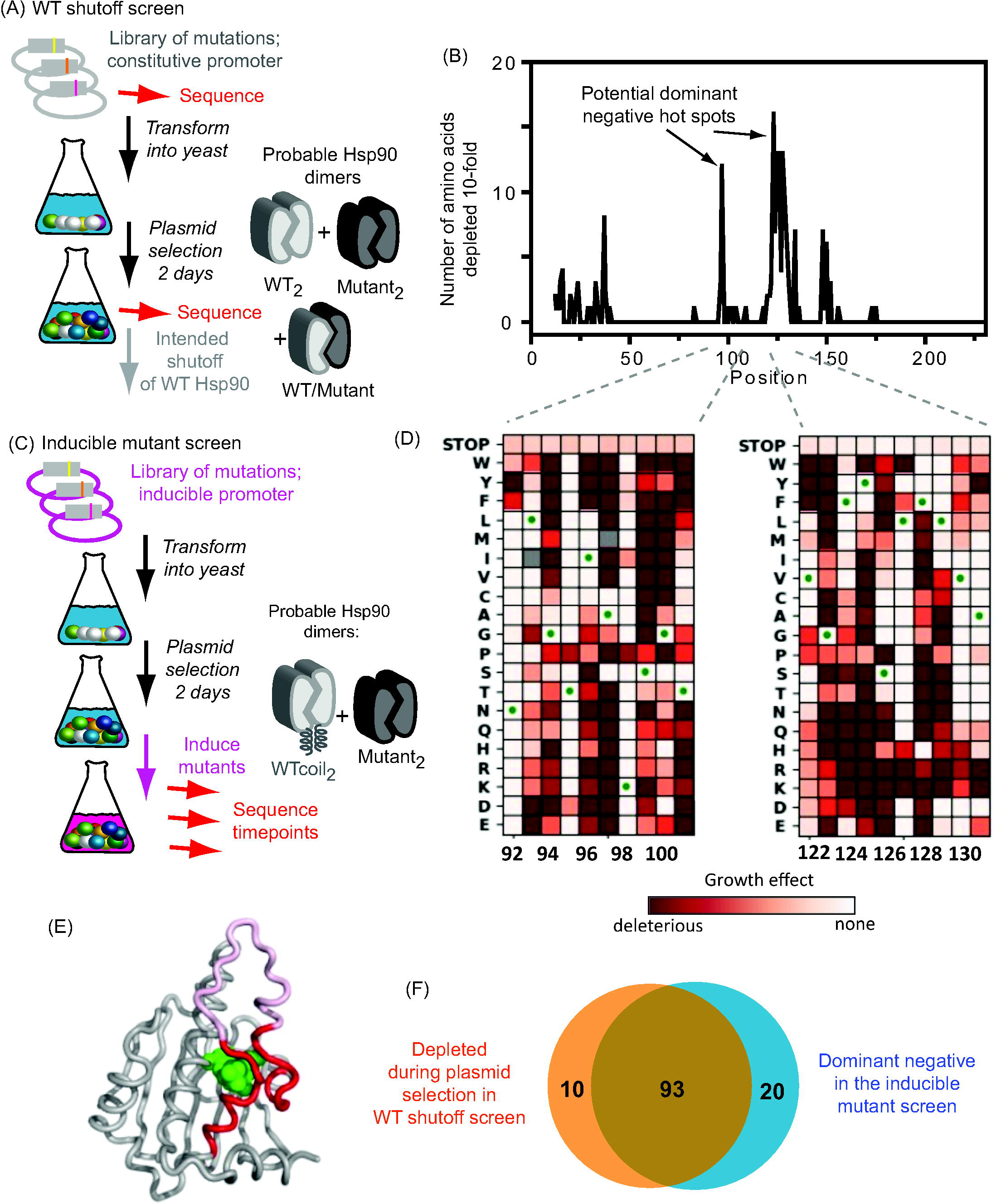
Comparison of screens for loss of function and dominant negative Hsp90 variants. (A) Experimental outline of the WT shutoff screen. (B) Number of amino acids that were more than 10-fold depleted after plasmid selection. The first 11 amino acids of Hsp90 were not screened because of failures in library construction. (C) Experimental outline of inducible mutant screen used to quantify the dominant growth effects of all point mutations in hot spot regions (positions 92-101 and 122-131). (D) Heatmap of dominant growth effects quantified in the inducible mutant screen. (E) Model of the N domain of Hsp90 with ATP bound (shown in green) based on 2CG9.PDB^24^. The hot spot regions are colored red and the rest of the lid pink. (F) Venn diagram indicating overlap between mutations identified as depleted during plasmid selection in the WT shutoff screen and dominant negative in the inducible mutant screen. The overlap is significant (p<0.00001, chi^2^).

The mutations that were strongly depleted during plasmid selection clustered in two hot spots that form hinge regions that mediate closing and opening of the lid that closes over ATP (Fig. 2B). The structure of Hsp90 undergoes large rearrangements of its three domains in response to ATP binding and hydrolysis^25^. As the lid region closes over and directly contacts ATP, conformational changes in the lid in response to ATP hydrolysis are thought to trigger the large rearrangement of domains from the ATP state to the ADP state of Hsp90^25,26^. The concentration of dominant negative mutations in the lid indicates that many perturbations to its dynamic conformational properties can lead to toxic effects. We focused our subsequent efforts on understanding the mechanisms of dominant negative mutations in the lid.

We developed an approach to quantify dominant effects in the regions that were hot spots for mutant depletion (Fig. 2B) while controlling cross-dimerization with co-expressed wild-type Hsp90 (Fig. 2C). The C-terminal domain of Hsp90 forms a stable dimer^30^ such that variants with mutations in the ATPase domain should form a distribution of mutant/wild-type heterodimers and mutant/mutant homodimers. To limit cross-dimerization (Fig. 2C), we utilized a yeast strain where the only chromosomal copy of Hsp90 has an appended coiled-coil that hinders it from associating with Hsp90 variants lacking the coiled-coil^31^.

To experimentally identify dominant negative Hsp90 mutations we drove expression of mutant variants in the lid hinge regions (aa 92-101 and 122-131) from the Gal1 promoter (Fig. 2C). The Gal1 promoter (pGal1) is tightly regulated so that there is no measurable expression when yeast are grown in dextrose, but strong expression is induced with galactose as the sugar source. Libraries of plasmid-encoded Hsp90 variants were transformed into yeast and the bulk culture expanded in dextrose media. Without expression of the mutant Hsp90, all variants could amplify independent of potential toxic effects of the encoded protein. Transfer of the bulk culture to galactose media then induced expression of the mutant Hsp90 variants along with WT Hsp90 whose expression continued to be driven by a constitutive promoter. Following induction with galactose, we collected samples at different time points and used deep sequencing to estimate the change in frequency of each mutant over time.

In the inducible mutant screen, there was a broad range of dominant growth effects (Fig. 3A and Table S2). There was a narrow clustering of effects for codons that encode the same amino acid (Fig. 3B), indicating both the reproducibility of the experiments and that selection was predominantly acting on the protein sequence and not the nucleic acid sequence. We also cloned individual dominant negative variants and confirmed their toxic effects for yeast growth both on plates and in liquid culture (Fig. 3C&D). We chose a conservative approach to categorize the strongest dominant negative substitutions with growth effects of less than -0.4, which represents four standard deviations less than the average stop codon. Of note, the toxic effects of stop codons are similar in magnitude to effects previously observed in yeast for the expression of folding compromised GFP^32^. Given the position of the lid region in the middle of the primary sequence of the N domain, stop mutations likely result in misfolded proteins that act to slow growth. Using the conservative cutoff, we identified 113 strongly dominant negative mutations in the hinge regions of the lid.

**Figure 3.**
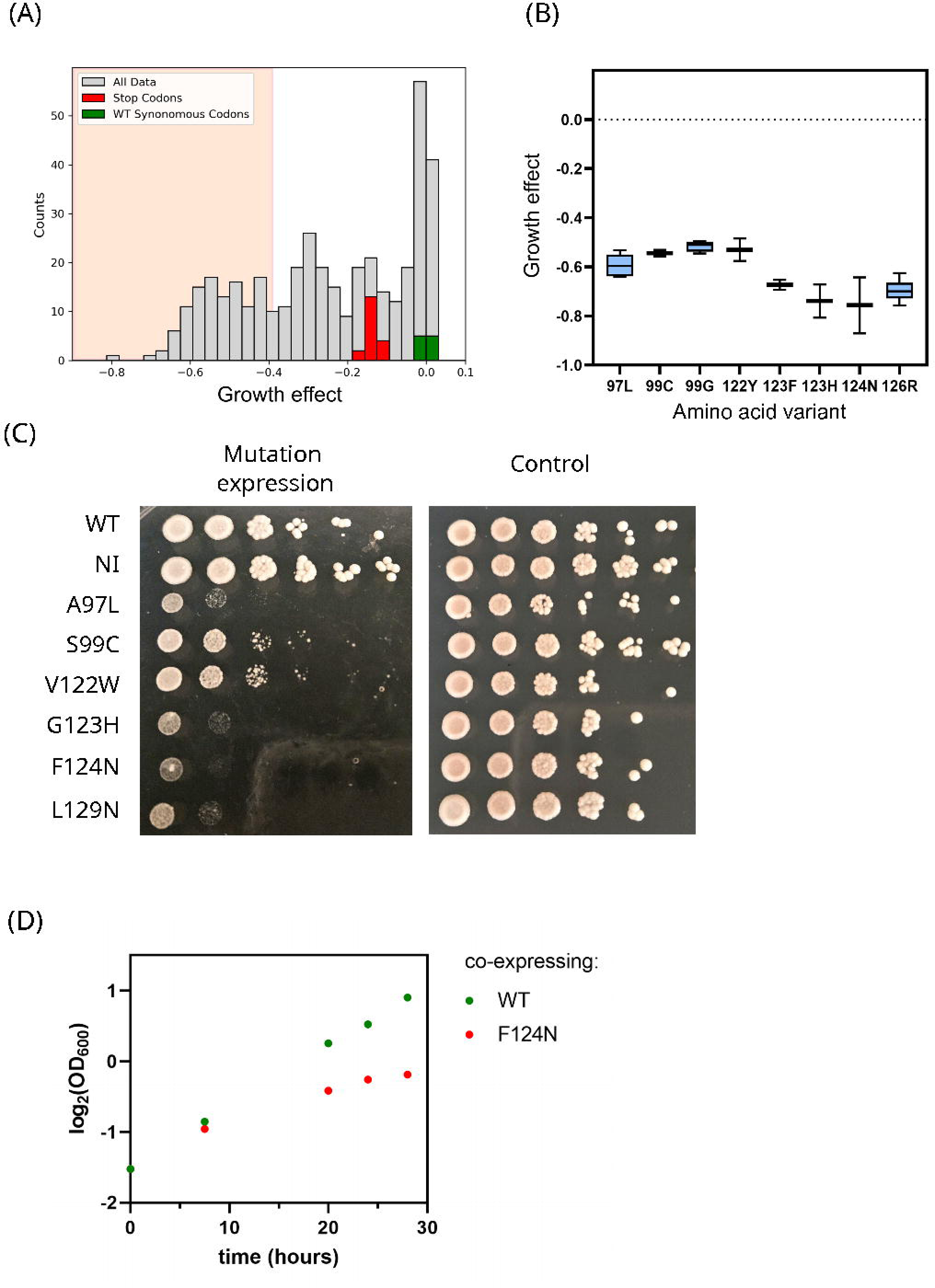
Reproducibility and validation of dominant negative variants. (A) Distribution of growth effects from the inducible mutant screen. Variants with growth effects lower than -0.4 were classified as dominant negative. (B) Variation in growth effects among synonymous codons for the same amino acid substitution. Growth properties of individually isolated dominant negative variants on plates (C) and in liquid culture (D).

There was large and significant (p < 0.0001 based on Chi^2^ test) overlap between the set of dominant negative mutations we directly observed in the hinge regions and mutations that were depleted during plasmid selection in the WT shutoff screen (Fig. 3F). This observation indicates that missing amino acids during plasmid selection in the initial screen were commonly due to dominant negative effects and not due to an inability of the plasmid-encoded variants to transform into cells. It also indicates that most dominant negative mutations have toxic effects independent of cross-dimerization with WT subunits because this cross-dimerization was limited in our induced mutant screen, but not our WT shutoff screen.

Motivated by the role of the lid region in closing over bound ATP, we examined how ATP hydrolysis impacted the dominant effects of individual mutations. We chose to investigate the A97L, G123H and F124N dominant negative mutations because they exhibited some of the strongest growth defects. To block ATP hydrolysis, we added the E33A mutation. In WT Hsp90, E33 positions a water molecule for ATP hydrolysis^24^. Previous studies have shown that the E33A mutation is compatible with ATP binding but prevents hydrolysis^33^. In the context of all three different dominant negative mutations we analyzed, E33A dramatically reduced toxic effects on growth (Fig. 4A). This result indicates that these mutations exert their toxic effects through a dynamic ATPase driven mechanism. Of note, pull down experiments of dominant negative variants from Hsp90 did not reveal any strong binding partners (Fig. S1), consistent with dynamic conformations and complexes leading to the observed toxicity.

**Figure 4.**
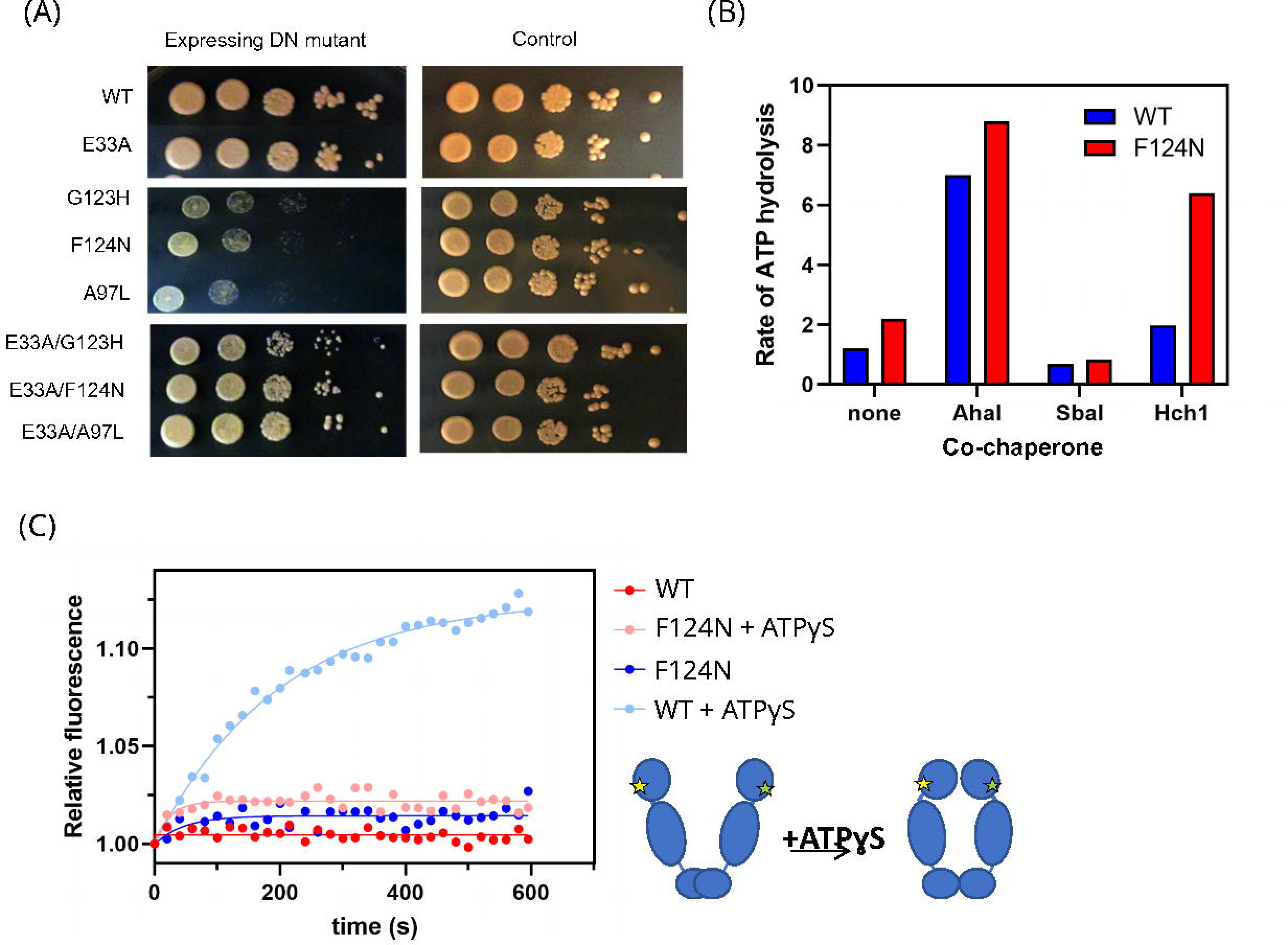
Impacts of dominant negative mutations on ATP-driven conformational changes. (A) The E33A mutation that prevents ATP hydrolysis in Hsp90 rescues the toxic effects of multiple dominant negative mutations. (B) ATPase rate of purified Hsp90 variants in the presence and absence of co-chaperones. (C) Conformation of fluorescently labeled WT and F124N Hsp90 protein monitored by FRET.

We further interrogated the role of ATPase activity on the biochemical properties of the F124N Hsp90 variant in purified form. F124N showed moderately elevated basal ATPase activity in vitro (Fig. 4B). The ATPase activity of F124N responded to co-chaperones similarly to WT Hsp90 – stimulated by Aha1 and Hch1 and retarded by Sba1. Previous studies have demonstrated that a wide range of at least two orders of magnitude of basal ATPase activity is compatible with robust yeast growth^34^. In purified form, the ATPase activity of F124N is within a range that is compatible with robust yeast growth and it responds to co-chaperones similarly to WT Hsp90. The toxic properties of F124N do not appear to be due to gross disturbance of ATPase activity, or co-chaperone interactions.

ATP binding induces a large open to closed conformational change in WT Hsp90^25^, and we examined how the F124N mutation impacted ATP-driven structural rearrangements (Fig. 4C). We used a FRET approach originally developed by Buchner and colleagues^35^ to monitor the conformation of purified full-length Hsp90 in the presence and absence of the ATP analogue ATPγS. Similar to previous observations, the ATP analogue stimulates a closed state for WT Hsp90. However, the FRET signal in the presence of ATPγS is greatly reduced for F124N compared to WT Hsp90, suggesting that F124N is not able to readily form a closed structure. Multiple studies including single molecule fluorescent analyses, indicate that the formation of a closed ATP bound structure is a consistent part of the ATPase driven conformational cycle of Hsp90^36^. Our findings with F124N indicate that ATPase activity does not require forming a closed conformation, but when the linkage between ATPase activity and formation of the closed structure is disrupted, it causes critical challenges for cell growth.

Motivated by the role of Hsp90 in triage decisions targeting clients for degradation in mammalian systems^17,19^ and the dynamic ATPase properties of F124N, we investigated client stability to proteolysis (Fig. 5). Induction of two different dominant negative Hsp90 variants (F124N and G123H) both led to large decreases in the levels of the model client protein GR (Fig. 5A). We also found that induction of F124N or G123H reduced expression of a GR-dependent reporter (Fig. 5B), consistent with the reduced expression of GR caused by the DN Hsp90 variants. The expression of GR in the presence of DN Hsp90 was rescued by the proteosome inhibitor MG132 (Fig. 5C&D). Together, these results demonstrate that dominant negative Hsp90 variants can lead to the proteasome-dependent degradation of a client protein.

**Figure 5.**
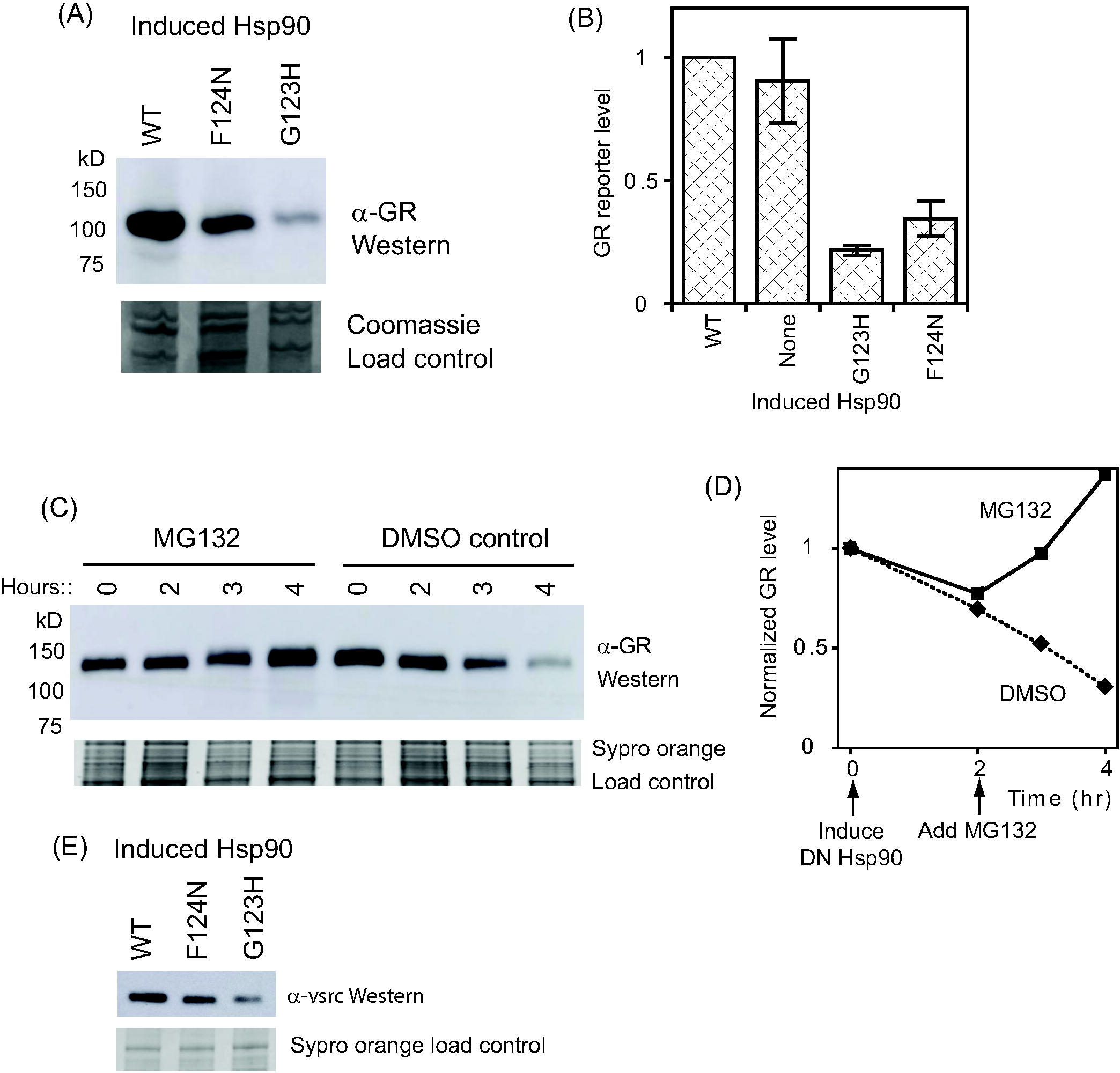
Dominant negative Hsp90 destabilizes a model client to degradation. Western blot analyses of steady state abundance of the model Hsp90 client GR in yeast expressing additional WT, F124N or G123H Hsp90. (B) GR-dependent reporter activity in yeast expressing additional WT or DN Hsp90 variants. Error bars represent the variation from two experimental replicates. (C) MG132 rescues expression of GR in yeast cells expressing G123H Hsp90. (D) Quantification of GR level from panel C. (E) The expression level of a second Hsp90 client (v-src) is reduced when F124N or G123H Hsp90 is induced in yeast.

Expression of F124N or G123H also resulted in decreased expression of a second model Hsp90 client, the v-src kinase (Fig. 5E). Of note, GR and v-src are structurally distinct clients that differ in the stringency that they rely on Hsp90 function^37^. The similar impacts of DN Hsp90 mutations on v-src and GR expression indicates that degradation is not client-specific and may be a general outcome for many clients.

Based on all our observations, we propose a model where formation of an ATP-bound closed Hsp90 structure is important for promoting the maturation of client proteins and dominant negative mutations target clients for proteasome degradation by hydrolyzing ATP from a more open structure (Fig. 6). This model is consistent with dominant negative mutant effects depending on ATP hydrolysis, failing to form a closed structure, and promoting the proteasome dependent destruction of a client protein.

**Figure 6.**
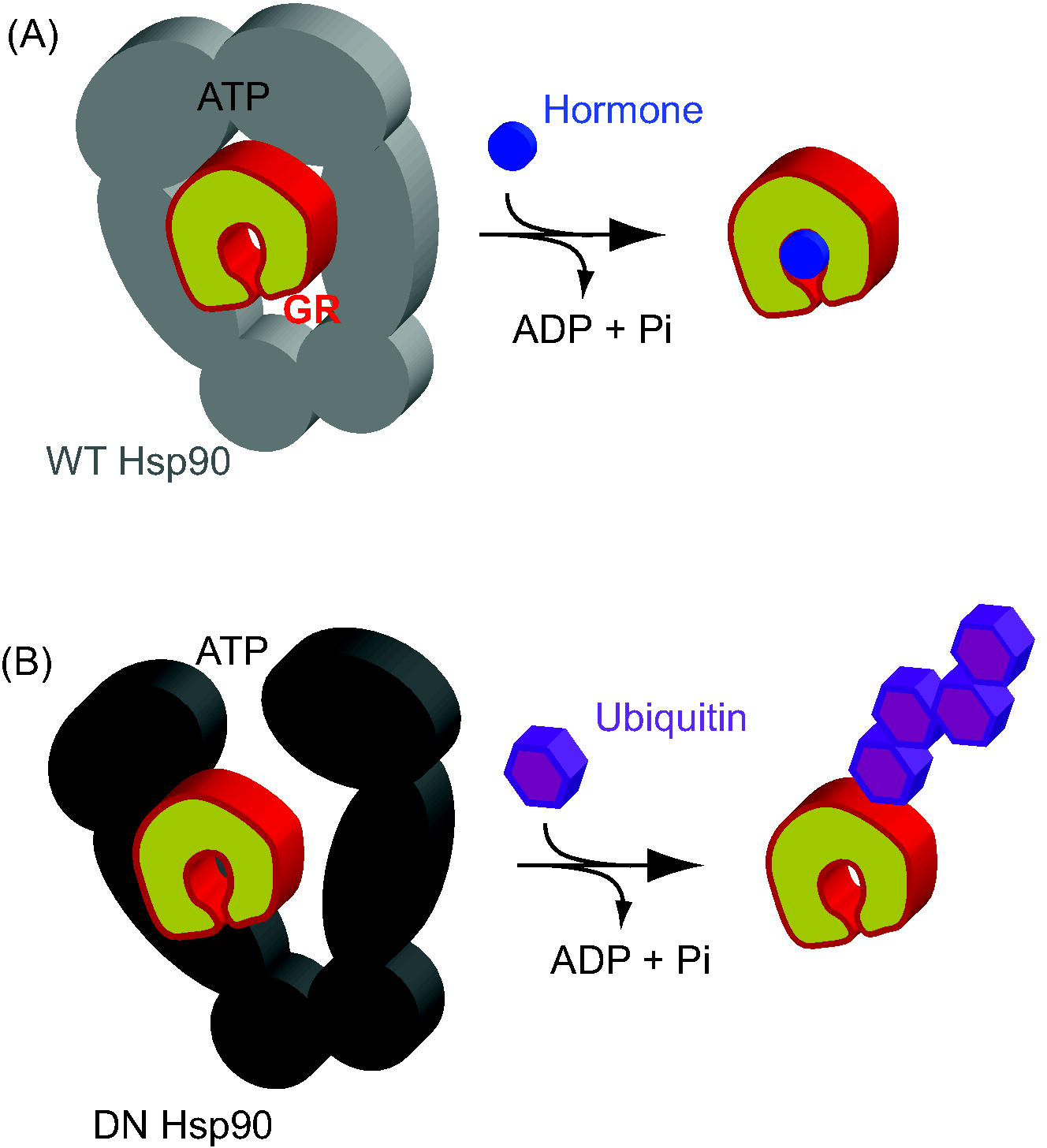
Model of the mechanism of dominant negative Hsp90 variants. (A) ATP hydrolysis in WT Hsp90 occurs in a fully closed state that protects client from being tagged for degradation. (B) In DN Hsp90, ATP hydrolysis can occur in a partly open structure that facilitates tagging of client for degradation.

## Conclusions

Our findings provide key evidence for an evolutionarily conserved function of Hsp90 in triage decisions for client proteins and provides insights into its mechanism. Prior studies had shown that the mammalian-specific Hsp90 co-chaperone CHIP can help to mediate triage decisions by Hsp90^18^. Multiple studies have also shown that competitive ATP inhibitors of Hsp90 lead to the ubiquitin targeted degradation of many clients in mammalian cells^17,20^. It has also been shown that many mammalian E3’s physically bind to Hsp90^10^, though it was unclear if this indicated that they were clients or acting to assist in triage decisions. Our work demonstrates that Hsp90 can target clients for degradation in yeast when ATP hydrolysis occurs without forming a fully closed state. This mechanism has several important potential implications that will be interesting to investigate in future studies. For example, we speculate that this mechanism of targeting clients for degradation may occur infrequently in the ATPase cycle of WT Hsp90 such that clients that stay bound through multiple cycles will eventually be targeted for degradation. In addition, this mechanism may be at play in the action of Hsp90 while bound to competitive ATP inhibitors. Our study demonstrates that triage decisions are an inherent and evolutionarily conserved property of Hsp90 and that hydrolysis of ATP in the closed state of Hsp90 is critical for keeping clients on the maturation pathway as opposed to the degradation pathway.

This work also demonstrates that dominant negative mutations can provide powerful new views of biologically important mechanism, especially for highly dynamic protein hubs such as Hsp90. Systematic identification of dominant negative mutations can be generally applied and used to study the mechanism of other important genes.

## Material and Methods

### Library construction

Point mutation libraries were generated in p417 plasmids using a cassette ligation strategy as previously described^38^. To construct the libraries for the inducible mutant screen, a destination vector was generated using a pRS414 plasmid containing the GalS promoter^39^ and the first and last 30 bases of Hsp90 bracketing a SphI restriction site. Two 10 consecutive amino acid saturation mutagenesis libraries covering amino acids 92-101 and 122-131 of Hsp90 were excised from the p417 plasmids using restriction enzymes that cut immediately upstream and downstream of the Hsp90 gene and annealed into the destination vector cut with SphI using SLIC cloning as described^29,40^. The number of independent transformants was in large excess to the library diversity.

### Strain construction

To avoid forming heterodimers between the mutant and wild-type versions of Hsp90, a strain was created in which a coiled-coil version of Hsp90 was the sole Hsp90 expressed in the yeast strain. To do this, we used the haploid Saccharomyces cerevisiae ECU82a plasmid swap strain, which is a derivative of W303 in which both endogenous Hsp90 genes, hsp82 and hsc82 are knocked out and wild-type hsp82 gene is expressed from a pKAT6 URA3 marked plasmid amenable to negative selection^28^. The Hsp90-coil construct containing a coiled-coil sequence (GGGTSSVKELEDKNEELLSEIAHLKNEVARLKKLVGERTG) inserted after amino acid 678 of Hsp90^31^ driven by a constitutive GPD promoter together with a HIS3 marker was integrated into the HO genomic locus of the plasmid swap strain. The resulting transformants were grown on plates containing 5-fluoroorotic acid, which selects for loss of the pKAT6 plasmid, to create the strain JFY12 (can1-100 ade2-1 his3-11,15 leu2-3,12 trp1-1 ura3-1 hsp82::leu2 hsc82::leu2 ho::pgpd-hsp82-coil-his3).

### Bulk competition experiment (Inducible mutant screen)

The JFY12 strain was transformed with the two p414Gal Hsp90 libraries (aa 91-101 and aa 121-131) using the lithium acetate method at a transformation efficiency sufficient to obtain a greater than 50-fold coverage of each mutant in each library (100,000 total transformants). Following 12 hours of recovery in synthetic dextrose lacking histidine (SD-H), transformed cells were washed three times in synthetic dextrose lacking histidine and tryptophan (SD-H-W to select for the presence of the Hsp90 library plasmid) to remove extracellular DNA, and grown in 50 mL SD-H-W media at 30°C for 24 hours with repeated dilutions to maintain the cells in log phase of growth and to expand the library. At least 10^7^ cells were passed for each dilution to avoid population bottlenecks. Cells in log growth were washed twice in synthetic 2% raffinose media lacking histidine and tryptophan (SRaf-H-W), diluted to early log phase in 50 mL SRaf-H-W and grown for 16 hours until the cultures reached mid-log phase. To begin the bulk competition experiment, cells were diluted to early log phase in synthetic 1% raffinose, 1% galactose media lacking tryptophan and histidine (SRafGal-H-W). An initial 0 hour time point was collected of ∼10^8^ cells, pelleted, washed, and stored at -80C. The remaining cells were grown for 30 hours at 30°C with samples collected as before at various time points (5, 8, 11, 14, 17, 25, 28 and 30 hours).

### Bulk competition experiment (WT shutoff screen)

For the single-expression competition experiment, point mutation libraries covering amino acids 12-231 were transferred as above to a p414GPD destination vector, which constitutively expresses Hsp90 at endogenous yeast levels ^28^. The libraries were transferred in 10 amino acid segments to ensure complete coverage of the library. The p414GPD Hsp90 libraries were then transformed into the DBY288 Hsp90 shutoff strain (can1-100 ade2-1 his3-11,15 leu2-3,12 trp1-1 ura3-1 hsp82::leu2 hsc82::leu2 ho::pgals-hsp82-his3)^41^ which was generated from the Ecu Hsp90 plasmid swap strain ^28^ by integration of Hsp90 driven by a Gal promoter^39^ together with a HIS3 marker into the HO genomic locus. Bulk competition experiments were performed as described^37^. In short, mutant libraries were amplified in the presence of wild-type Hsp90 in SRafGal-H-W, and then switched to wild-type shutoff conditions in SD-H-W media for 16 hours at 30°C. A sample of the culture was pelleted and stored at -80°C at this point and after a further 8 hours of competition. Libraries containing 10 consecutive amino acids were competed and analyzed in separate cultures to facilitate efficient analyses of mutant frequency using focused next-generation sequencing. All mutants were analyzed in parallel in the same batch of media with competition cultures grown on the same day in the same incubator. Cultures were maintained in log phase by regular dilution with fresh media and managed to maintain a population size of 10^9^ or greater to minimize impacts from bottlenecking. Sample processing for next-generation sequencing and data analyses were performed as previously described^38^.

### Yeast growth analysis

To determine the growth rate of individually cloned Hsp90 mutants, a panel of Hsp90 point mutations were individually cloned by site-directed mutagenesis into the p414Gal vector and transformed into JFY12 using a lithium-acetate based procedure. Individual clones were selected on SD-H-W media where expression of the mutants was not induced. Clones were grown in liquid SD-H-W media to a cell density of 10^8^ at 30°C, washed with SRaf-W-H to remove residual dextrose and diluted to early log growth phase in SRaf-H-W and grown for 16 hours at 30°C. Cells were then diluted to early log phase in SRafGal-H-W to induce the expression of the mutants. The growth rate of the cultures was monitored based on absorbance at 600 nm over time and fit to an exponential growth model ^29^. During this time, the cells were maintained in log phase by periodic dilution. Additionally, after 8 hours of growth in the SRafGal-H-W induction media, spotting assays were performed. Serial 10-fold dilutions of each culture was spotted on both SRafGal-W-H (to induce mutant Hsp90 expression) and SD-H-W (control – no mutant expression) and incubated for 2 days at 30°C to monitor colony growth.

### Analysis of steady state levels of GR and v-src and measurement of GR activity

To determine the impact of the DN Hsp90 variants on the steady state levels of GR and v-src, the JFY12 strain was transformed with p414Gal plasmids expressing either WT, F124N or G123H Hsp90 and either the P2A/GRGZ plasmid described previously that constitutively expresses GR^42^ or the p316Gal-vsrc-V5 plasmid described previously^23^ that expresses v-src upon addition of galactose. Cells were grown as above and harvested after 6 hours of growth in inductive media (SRafGal). Frozen yeast pellets were thawed on ice and lysed using glass beads in a lysis buffer containing 5mM EDTA, 50mM Tris pH 8, and both 5mM PMSF and protease inhibitor cocktail as protease inhibitors. The total protein concentration in each lysate was estimated using a BCA assay. The lysate was analyzed via SDS-PAGE electrophoresis and western blotting. The PVDF membrane was blocked in 5% milk and TBS (tris buffered saline) at room temperature for 1 hour. After blocking, the membrane was either probed with the FLAG tag primary antibody for GR (FG4R, Invitrogen), or the V5 tag primary antibody for v-src (V4RR, Invitrogen) at a 1:5000 dilution in 1% milk and TBS overnight at 4°C. After washing in 1xTBS, the membrane was incubated in anti-mouse secondary conjugated to HRP at a 1:5000 dilution in 4% milk and TBS for 1 hour at room temperature. The surface of the membrane was incubated in ECL substrate for 5 minutes. Bands were visualized using chemiluminescence. β-galactosidase reporter activity of yeast lysates was quantified as described previously^23^.

### Analysis of steady state levels of GR following proteosome inhibition

To measure the stability of GR in the presence and absence of the proteosome inhibitor MG132, experiments were performed as above with the following modifications. To increase uptake of MG132 in our yeast strains L-proline was used instead of ammonium sulfate in all growth media as the sole nitrogen source and a small amount of sodium dodecyl sulfate (SDS) was added prior to MG132 treatment as previously described^43^. These treatments are thought to lead to a transient opening of the cell wall/membrane. At the point at which cells were switched to SRafGal-W-A media to induce expression of the mutant proteins, 0.003% SDS was added and cells were then grown at 30°C for two hours. At this point, either 50 µM MG132 (Sigma, M7449) or an equivalent volume of DMSO as a control was added. Cells were then grown at 30°C and samples were harvested and frozen at -80°C at 0, 1 and 2 hours following treatment.

### Purification of Hsp90

Hsp90 variants with N-terminal His_6_ tags were bacterially expressed, purified and analyzed as described previously ^23^. In brief, protein purification was performed using Co+NTA agarose (Qiagen), followed by a Phenyl Sepharose column, and a Q sepharose HP column (GE). Aha1 and Hch1 were purified as previously described^42,44^. The concentrations of these highly purified proteins were determined spectroscopically using extinction coefficients based on amino acid composition.

### ATPase assay

Hydrolysis of ATP by Hsp90 was determined using an NADH-coupled ATPase assay as previously described^45^. ATPase assays were performed at 37°C in 20 mM Tris pH 7.5, 5 mM MgCl2, 100 mM KCl using a Bio50 Spectrophotometer equipped with a Peltier temperature control unit (Cary) and a 1 cm pathlength cuvette. The reaction was initiated by the addition of 2 mM ATP. Absorbance at 340 nm (ε340 = 6220 M^-1^ cm^-1^) was measured at 15 second intervals for 10 min as previously described^23^.

### FRET labeling and assay

Hsp90 was labeled with donor and acceptor dyes (ATTO-488-maleimide and ATT0-550-maleimide) (company) as recommended by the manufacturer at an engineered cysteine residue (D61C)^35^. 2 µM donor-labeled and 2 µM acceptor labeled Hsp90 were mixed in 20 mM Tris pH 7.5, 5 mM MgCl_2_, 100 mM KCl and incubated at 37°C for 30 minutes to allow subunit exchange followed by addition of the indicated concentration of ATP⍰S and FRET was measured in a Fluoromax 3 machine at 37°C.

### Hsp90 affinity isolation

Hsp90-proteinA affinity capture was performed as before with the following modifications^46,47^. A gene encoding protein A was inserted after amino acid position 684 of hsp82 and cloned into the p414Gal plasmid. The JFY12 stain was transformed with either p414Gal hsp82-proteinA (WT) or p414Gal hsp82-proteinA (F124N) plasmids. Cells were grown first in SD-H-W media at 30°C to stationary phase, diluted to early log phase in SRaf-H-W (2 L) and grown for 16 hours and then 1% galactose was added to the cultures to induce expression of the Hsp90 variants and grown for an additional 6 hours at 30°C. Cells were harvested, washed with water, pressed through a syringe into liquid nitrogen to make frozen noodles, and stored at -80°C. Frozen cells were then lysed by grinding in a Planetary Ball Mill PM 100 machine. 150 mg of the resulting cell powder was resuspended in 50 mM HEPES pH 7.5, 150 mM NaCl, 20 mM Na_2_MoO_4_, 2 mM EDTA, 5% glycerol, 1x protease inhibitor cocktail. The lysate was clarified by centrifugation at 13k rpm for 10 min at 4°C and the resulting supernatant was added to 20 µl of homemade rabbit IgG magnetic beads. Cell lysate was incubated with beads for 1 hour at 4°C to allow binding. Bound beads were quickly washed three times with cold lysis buffer and then resuspended in 1x SDS lysis buffer. Samples were run on SDS-PAGE gel and stained with Sypro Orange stain (Thermo Fisher Scientific).

## Acknowledgements

This works was supported by grant R01-GM112844 from the National Institutes of Health to D.N.A.B.

