## Supplementary material for "Dominant negative mutations in yeast Hsp90 reveal triage decision mechanism targeting client proteins for degradation": Figure S1

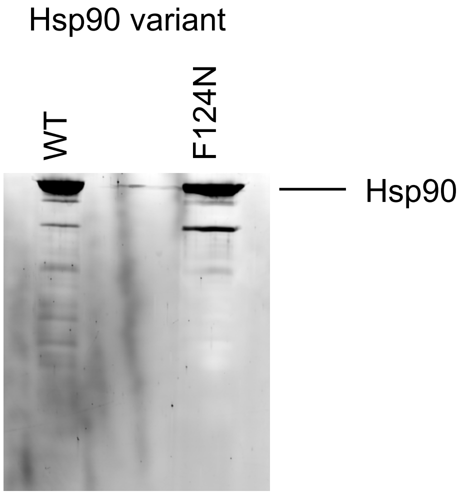


Figure S1. Pull-down assays were carried out using lysates from yeast cells expressing WT or F124N Hsp90 tagged with Protein A (see Materials and Methods). The proteins were visualized by sypro orange stain.
